# Three-dimensional random walk models of individual animal movement and their application to trap counts modelling

**DOI:** 10.1101/2020.07.28.224766

**Authors:** DA Ahmed, S Benhamou, MB Bonsall, SV Petrovskii

## Abstract

**Background:** Random walks (RWs) have proved to be a powerful modelling tool in ecology, particularly in the study of animal movement. An application of RW concerns trapping which is the predominant sampling method to date in insect ecology, invasive species, and agricultural pest management. A lot of research effort has been directed towards modelling ground-dwelling insects by simulating their movement in 2D, and computing pitfall trap counts, but comparatively very little for flying insects with 3D elevated traps.

**Methods:** We introduce the mathematics behind 3D RWs and present key metrics such as the mean squared displacement (MSD) and path sinuosity, which are already well known in 2D. We develop the mathematical theory behind the 3D correlated random walk (CRW) which involves short-term directional persistence and the 3D Biased random walk (BRW) which introduces a long-term directional bias in the movement so that there is an overall preferred movement direction. In this study, we consider three types of shape of 3D traps, which are commonly used in ecological field studies; a spheroidal trap, a cylindrical trap and a rectangular cuboidal trap. By simulating movement in 3D space, we investigated the effect of 3D trap shapes and sizes and of movement diffusion on trapping efficiency.

**Results:** We found that there is a non-linear dependence of trap counts on the trap surface area or volume, but the effect of volume appeared to be a simple consequence of changes in area. Nevertheless, there is a slight but clear hierarchy of trap shapes in terms of capture efficiency, with the spheroidal trap retaining more counts than a cylinder, followed by the cuboidal type for a given area. We also showed that there is no effect of short-term persistence when diffusion is kept constant, but trap counts significantly decrease with increasing diffusion.

**Conclusion:** Our results provide a better understanding of the interplay between the movement pattern, trap geometry and impacts on trapping efficiency, which leads to improved trap count interpretations, and more broadly, has implications for spatial ecology and population dynamics.

## 1 Introduction

Modelling individual animal movement and navigation strategies using random walks has long been a successful tradition in movement ecology (Nathan et al., 2008). The earliest models considered animal paths as uncorrelated and unbiased, e.g. Simple Random Walks (SRW) (Lin and Segel, 1974; Okubo, 1980). A natural extension known as the Correlated Random Walk (CRW), firstly conceived by Patlak (1953) and later developed by others (Hall, 1977; Kareiva and Shigesada, 1983; Bovet and Benhamou, 1988; Benhamou, 2004), allows for correlation between the orientations of successive steps, resulting in a short term localized directional bias known as ‘forward persistence’. This provides a more realistic description, as animals in the short term are more likely to keep moving in the same direction than to perform abrupt turns. Alternatively, a movement can show a consistent long term directional bias reflecting an overall preferred direction. This type of movement is known as a Biased Random Walk (BRW) (Marsh and Jones, 1988). If both short and long term biases are combined we obtain a Biased Correlated Random Walk (BCRW), (Benhamou, 2006; Codling et al., 2008; Bailey et al., 2018).

A tractable link between the 2D balanced CRW (i.e. left and right turns are equiprobable) and the mean squared displacement (MSD) was introduced by Tchen (1952) with constant step length, and later by Hall (1977) for variable step length. This helped bridge the gap between theory and field data, by providing a measure of the spatial spread of a population with the path length in terms of simple statistical moments. Kareiva and Shigesada (1983) further extended these results for a non-balanced 2D CRW. By comparing the observed MSD against that computed from theory, one could determine how well the CRW model predicted real animal movement (Weiss, 1994; Codling et al., 2008). This gave rise to a multitude of studies which successfully modelled the movement of a variety of species using the CRW, with many examples, including beetles (Byers, 2001), butterflies (Schultz and Crone, 2001), Elk (Morales et al., 2004; Fortin et al., 2005), grey seals (McClintock et al., 2012), and many others.

With cutting-edge developments in tagging and sensor technology, it is now possible to obtain accurate and refined 3D movement data, used to infer individual posture and heading (or 3D orientation). Measures of azimuthal, elevation and bank angles can be obtained through the usage of accelerometers and magnetometers, whereas, gyrometers can provide direct measures of rotations such as yaw, pitch and roll (Williams et al., 2020). Alongside this, there has been an increase in the number of studies which focus on 3D animal movements (Voesenek et al., 2016; Le Bras et al., 2017; de Margerie et al., 2018). In light of the above context, an extension to the results conceived by Hall (1977) to 3D is evidently due. Recently, Benhamou (2018) derived a mathematical expression for a key metric, namely, the MSD of the balanced CRW in 3D space (which can easily be extended to BRWs), and also path sinuosity, which is directly linked to the MSD of CRWs and expresses the amount of turning associated with a given path length. This sets the stage for 3D CRWs and 3D BRWs to be tested as null models that could hypothetically provide a more realistic framework for swimming, burrowing and flying animals – due to the mere fact that movement is exercised in an additional (third) direction. Once the above movement models are formalised, these can then be used as a baseline for a theoretical insight into the dynamics of trap counts.

Trapping is the predominant sampling method in insect ecology, invasive species, and agricultural pest management. Their usage covers a wide scope of ecological scenarios, including; general survey of insect diversity, detection of new invasive pests, delimitation of area of infestation, suppressing population buildup, monitoring populations of established pests, or even as a direct control measure, etc. (Southwood, 1978; Radcliffe et al., 2008). Since their original conception, many traps have been designed with modifications to cater for particular species, habitats, and research requirements (Muirhead-Thomson, 1991). Considerable progress has been made in modelling 2D pitfall trapping systems (Petrovskii et al., 2014), with recent efforts to standardize methodology (Brown and Matthews, 2016), however, few attempts can be found in the literature which analyse 3D elevated traps, albeit some efforts entirely based on simulations (Byers, 2011, 2012). We are interested in those traps used for flying insects. For this purpose, the main two types which are used in ecological studies are the ‘interception’ trap in the form of a net-like structure e.g. Malaise trap (tent-shaped; Lamarre et al., 2012), or ‘sticky’ traps usually coated with an adhesive. We focus on the latter, which, from a mathematical perspective, constitutes an enclosed shape with absorbing surface. In agricultural studies, the most commonly used traps are sticky spheroidal, cylindrical, and cuboidal traps, particularly for faunal surveys (Taylor, 1962; Sivinski, 1990; Robacker and Rodriguez, 2004; Epsky et al., 2004). Amongst these, the default choice is usually the sticky spherical trap, which is known to effectively trap a variety of taxa, e.g. *Tephritid* fruit flies, such as; apple maggot flies (*Rhagoletis pomonella*), blueberry maggot flies (*Rhagoletis mendax*), papaya fruit flies (*Toxotrypana curvicauda Gerstaecker*) and biting flies in the family *Tabanidae* (Sivinski, 1990; Duan and Prokopy, 1994; Mondor, 1995; Kirkpatrick et al., 2017). It is also worth mentioning that other trap types do exist, but are used less frequently, for e.g. triangle (or wedge), diamond, cones and some others (Epsky et al., 2004), but usage largely depends on the target species.

In this paper, we provide the mathematical details behind modelling individual animal movement using a 3D SRW, and demonstrate how short/long term persistence mechanisms can be incorporated, for a more general and realistic 3D CRW or 3D BRW. Using the results from Benhamou (2018), we summarize important metrics, such as the MSD, and show how these RWs can be made equivalent in terms of diffusion. Using this 3D RW framework, we model the movement of animals in 3D space, with focus on trapping. We reveal that trap counts vary non-linearly as a function of trap surface area or volume, and provide analytic expressions useful for trap count estimation. Furthermore, we investigate the interplay between the trap shape and elongation of 3D traps, the movement behaviour and how this can induce changes in trapping efficiency. More specifically, we analyse the impact of trap geometry and how short-term correlations (‘micro-structure’) or diffusion (‘macro-structure’) can affect capture rates. Better understanding of trap count dynamics and catch patterns lead to improved trap count interpretations. More generally, the implications of our results are also relevant in other ecological contexts, for e.g. where trap size can be thought of as odour plume reach (Miller et al., 2015).

## 2 Methods

### 2.1 3D Random Walks in Cartesian and Spherical co-ordinates

Individual animal movement can be modelled in 3D as a time series of locations **x**_*i*_ = (*x*_*i*_, *y*_*i*_, *z*_*i*_) recorded at discrete times *t*_*i*_ = {*t*_0_,*t*_1_,*t*_2_, …}. The movement can therefore be seen as a series of discrete steps Δ**x**_*i*_ = **x**_*i*_ − **x**_*i*−1_. Any 3D RW can be described in spherical coordinates, by expressing the step vector in terms of step lengths *l*_*i*_ = ‖Δ**x**_*i*_‖, azimuthal angle *θ* _*i*_ (equivalent to longitude) and polar angle *ϕ* _*i*_ (equivalent to co-latitude), using the transformation:

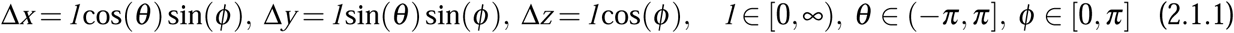

The change of direction of an animal from heading (*θ* _*i*_, *ϕ* _*i*_), between locations **x**_*i*−1_ and **x**_*i*_, to heading (*θ* _*i*+1_, *ϕ* _*i*+1_), between locations **x**_*i*_ and **x**_*i*+1_, can be modelled as an orthodromic (or great circle) arc, characterized by two angles: the initial arc orientation *β* _*i*_, measured between −*π* and *π* in the frontal plane with respect to the horizontal level, and the arc size *ω* _*i*_, measured between 0 and *π* in the plane defined by the two headings:

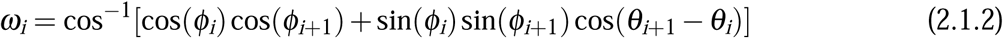

For a balanced CRW (including SRW as a special case) or BRW, the random variable *β* is independent of *ω*, and its distribution must also be centrally symmetric so that its mean sine and cosine are both null. Whether short or long term directional persistence is incorporated into the RW can be realised through the mean cosine of *ω, c*_*ω*_: one gets *c*_*ω*_ > 0 for a balanced CRW and BRW and *c*_*ω*_ = 0 for a SRW. CRW and BRW can be further distinguished based on how the heading at any step is determined. For both types of walks it is drawn at random around a predefined 3D direction µ. For a CRW, µ corresponds to the heading at the previous step, whereas for a BRW, µ corresponds to the target direction. In this case, the arc size corresponding to the angular discrepancy between a given heading and the target direction will be referred to as ν, which is statistically related to the arc size between successive headings *ω* through the relationship 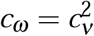 as occurs with 2D BRW (Marsh and Jones, 1988; Benhamou, 2006; Codling et al., 2008).

The Mean Squared Displacement (MSD), 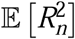, which is defined as the expected value of the squared beeline distance between an animals’ initial and final locations after *n* steps, serves as a useful metric to analyse movement patterns. The general MSD formulation for 2D CRW (Kareiva and Shigesada, 1983; Benhamou, 2006), in which left and right turns are not necessarily balanced, is extremely complex. We will consider here its extension in 3D space only for balanced CRW, developed by Benhamou (2018), and which reads:

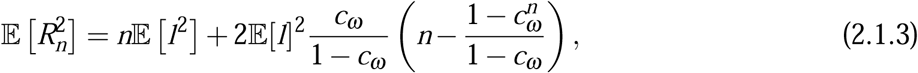

For a large step number *n*, the MSD approaches:

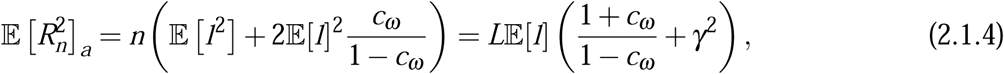

where *L* = *n*𝔼[*l*] is the mean path length and 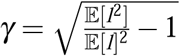 is the coefficient of variation of step length. For a 3D SRW, with *c*_*ω*_ = 0, the MSD reduces to 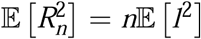, whatever the step number. It is readily seen from equation (2.1.4) that the MSD is asymptotically proportional to *n*, and therefore the walk becomes isotropically diffusive in the long term. The subscript ‘*a*’ is included here to represent the asymptotic value to which the MSD tends when *n* increases indefinitely. For an isotropically diffusive RW, the MSD is related to the diffusion coefficient *D* as follows: 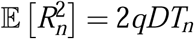 where *T*_*n*_ is the duration of the *n* step RW and *q* = 1, 2, 3 corresponds to the number of dimensions (Crank, 1975; Turchin, 1998; Sornette, 2004; Codling et al., 2008). The amount of turning in a random search path can be quantified by the sinuosity index:

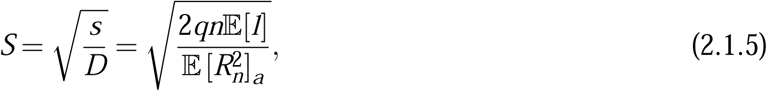

where *s* is the mean speed, with *q* = 3 for a random walk in 3D space (Benhamou, 2006, 2018).

In the case of a BRW, headings are drawn independently of each other in the target direction. This leads to the following expression for the MSD:

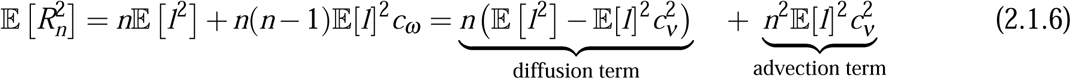

where *ν* is the arc size between an heading and the target direction, which is statistically related to the arc size between successive headings *ω* through the relationship 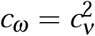. This expression highlights that a BRW is essentially a combination of the diffusive random walk and a drift, and its MSD is dominated in the long-term by the contribution of the drift. It is worth noting that the MSD expressions for balanced CRW and BRW in 3D space are similar to those obtained in 2D space (Hall, 1977; Marsh and Jones, 1988). The only difference is that the mean cosine of turning angles that is used in 2D space is replaced by the mean cosine of orthodromic arcs corresponding to the reorientations between successive 3D headings.

We can derive the conditions under which two 3D balanced CRWs are ‘equivalent’, in the sense that they have the same MSD after *n* steps, given that *n* is sufficiently large. In particular, if we consider a SRW with step length *l*^∗^ and mean cosine 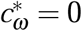, assuming the same coefficient of variation of step length and the same mean path length *L*, we obtain the following ‘condition of equivalence’:

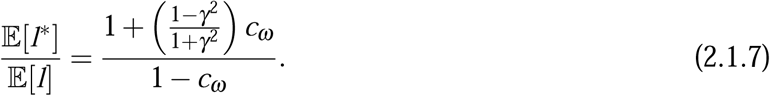

Now consider a SRW and a BRW with step lengths *l*^∗^ and *l*′ and mean cosines 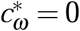 and 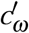, respectively. The condition of equivalence between these RWs in terms of diffusion is obtained with:

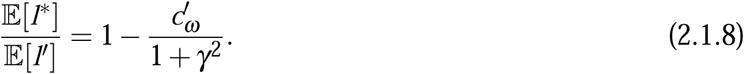

### 2.2 Mathematical bases for simulations of 3D RW

We relied on a distribution of step length so that the distributions of increments Δ*x*, Δ*y*, and Δ*z*, when reorientations are purely random (SRW), are zero-centred Gaussian distributions with the same standard-deviation *σ*, which represents the mobility of the animal. For a SRW with such increments, the probability that the animal moves into an (infinitesimally) small vicinity of the current location **x**, i.e within volume *dV* = *d*Δ*xd*Δ*yd*Δ*z*= *l*^2^ sin(*ϕ*)*dldθdϕ*, is:

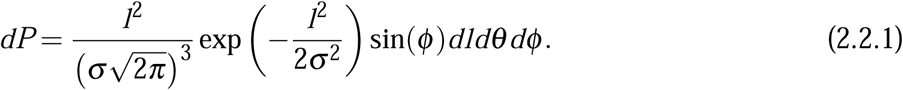

As *l, θ* and *ϕ* are mutually independent random variables, one gets the following probability distribution functions for these variables:

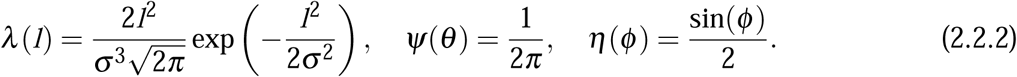

The mean step length is 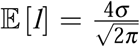 and mean squared step length 𝔼[*l*^2^] = 3*σ* ^2^. The coefficient of variation is therefore 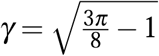. Note that, as expected, λ(*l*) can be considered a transformation of the Chi distribution with 3 degrees of freedom, for re-scaled step lengths 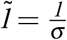 (Walck, 2007).

To specify the distributions of initial arc orientation *β* and arc size *ω* in our simulations, we used the von-Mises Fisher distribution (vMF), which is the simplest type amongst the Generalized Fisher-Bingham family of spherical distributions (Kent, 1982). The vMF distribution on the (*q* − 1)-dimensional sphere 𝕊^*q*−1^ in ℝ^*q*^ of the unit random vector **z** = (*z*_1_, *z*_2_, …, *z*_*q*_) is given by:

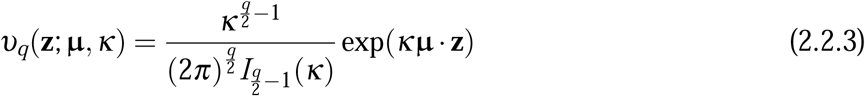

where µ is the mean direction with norm ‖µ‖ = 1 and *κ* > 0 is a measure of the concentration about the mean direction, and *I*_*m*_ denotes the modified Bessel function of the first kind of order *m*. For *q* = 2, this corresponds to a particular type of distribution on a circle, known as the von Mises distribution. For *q* = 3 on the sphere 𝕊^2^, with 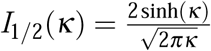 (Mardia et al., 1979), the probability density function of the endpoint of **z** falling within the infinitesimal surface element with surface area *ds* is:

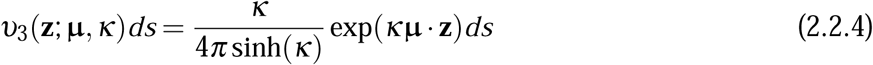

Given that µ and **z** are two unit vectors which deviate by *ζ* from each other, one gets µ · **z** = cos(*ζ*), with *ζ* = *ω* for a balanced 3D CRW, where µ corresponds to the previous heading, or *ζ* = ν for a 3D BRW, where µ corresponds to the target direction. Furthermore, by setting the pole of the sphere at the endpoint of µ, the infinitesimal surface element *ds* can be rewritten without loss of generality as sin(*ζ*)*dβdζ* as it appears that *ζ* then behaves as a co-latitude and *β* as a longitude. With *β* uniformly distributed between –*π* and *π*, one gets:

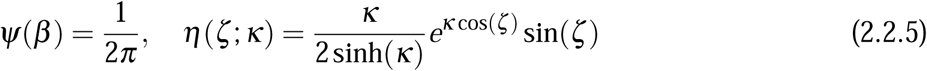

where *ψ* and η correspond to the probability distribution functions of the initial arc orientation and arc size, respectively (Fisher et al., 1981; Mardia and Jupp, 2000), with:

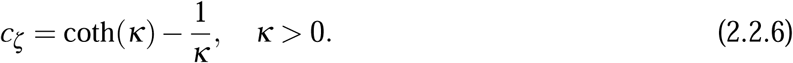

In the limit *κ* → 0, the distribution of the arc size simplifies to 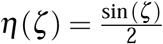 with mean cosine *cζ* = 0, as expected for a SRW.

**Figure 2.2.1.**
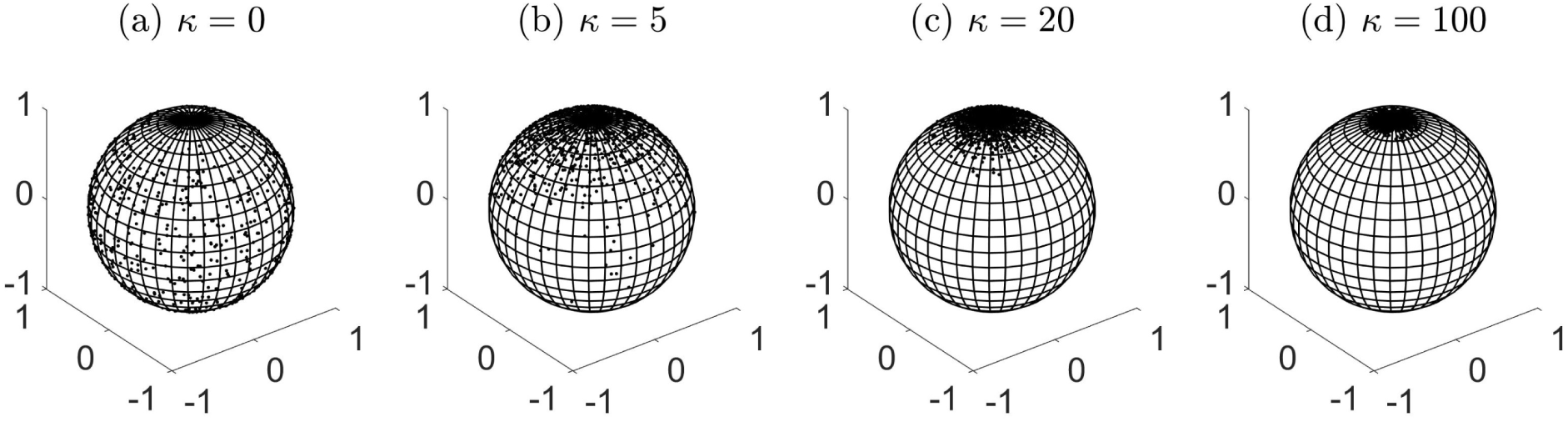
Random samples from the vMF distribution on a unit sphere 𝕊^2^ with the pole as the mean direction µ = (0, 0, 1), with increasing concentration parameter *κ*, based on 1000 simulated points.

In the case of a SRW, the points are uniformly distributed on the whole surface. For increasing *κ* values, the points are more concentrated towards the pole µ. Directional correlation is introduced to get a 3D balanced CRW, by randomly generating a heading from a distribution where the mean direction µ corresponds to the previous heading, whereas a 3D BRW is obtained by randomly generating a heading from a distribution where the mean direction µ corresponds to the target direction.

To tune the scale parameters of various CRW so that they are equivalent in terms of diffusion, we can express equation (2.1.7) as:

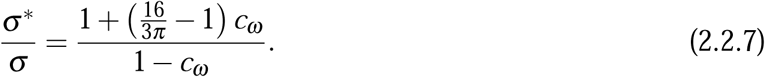

In the long term, a CRW with scale parameter *σ* behaves as a SRW with scale parameter *σ* ^∗^. The sinuosity of both walks can therefore be expressed as:

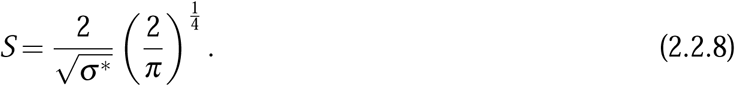

Similarly, for a BRW with step length distribution parameter *σ* ′ and long term persistence parameter *κ* ′ one gets:

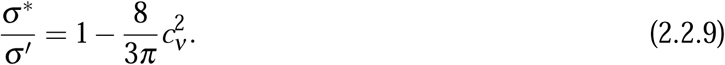

with 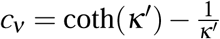.

### 2.3 Modelling trapping

In 3D trapping scenarios, consider a population of *N* individuals moving independently of each other. The path of each individual is modelled as a 3D RW in unbounded space, with initial location **x**_0_ = (*x*_0_, *y*_0_, *z*_0_) in proximity of a 3D trap. Each subsequent step is determined by the recurrence relation **x**_*i*_ = **x**_*i*−1_ + Δ**x**_*i*_, resulting in a RW which is governed by the type of probability distribution for the step vector (Δ**x**), and its properties. We assume that each walker moves until it is trapped or has travelled a path of length *L*, which can be easily converted into time by considering the mean speed *s*. We introduce the concept of trapping by stating that at each step *i*, any individual which is within the confines of a trap is removed from the system, leading to trap counts or captures. Under such conditions, the trap surface is absorbing and the simulation allows cumulative trap counts 𝔗 to be recorded. In our simulations, we assume the absence of mortality or reproduction, so that the population at each step can only decrease, due to trapping, or otherwise remains stable. As an example, Fig. 2.3.1 shows the distribution of the individuals over the 3D space after performing the random walk of a given length *L*.

In this study, we consider three shapes of 3D traps, namely the spheroid, cylindrical and rectangular cuboid types with trap geometry D defined by the following:

1. Spheroid (i.e. ellipsoid of revolution) trap with equatorial radius *r*_*s*_ and polar radius *h*_*s*_,

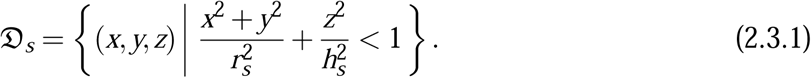

**Figure 2.3.1.**
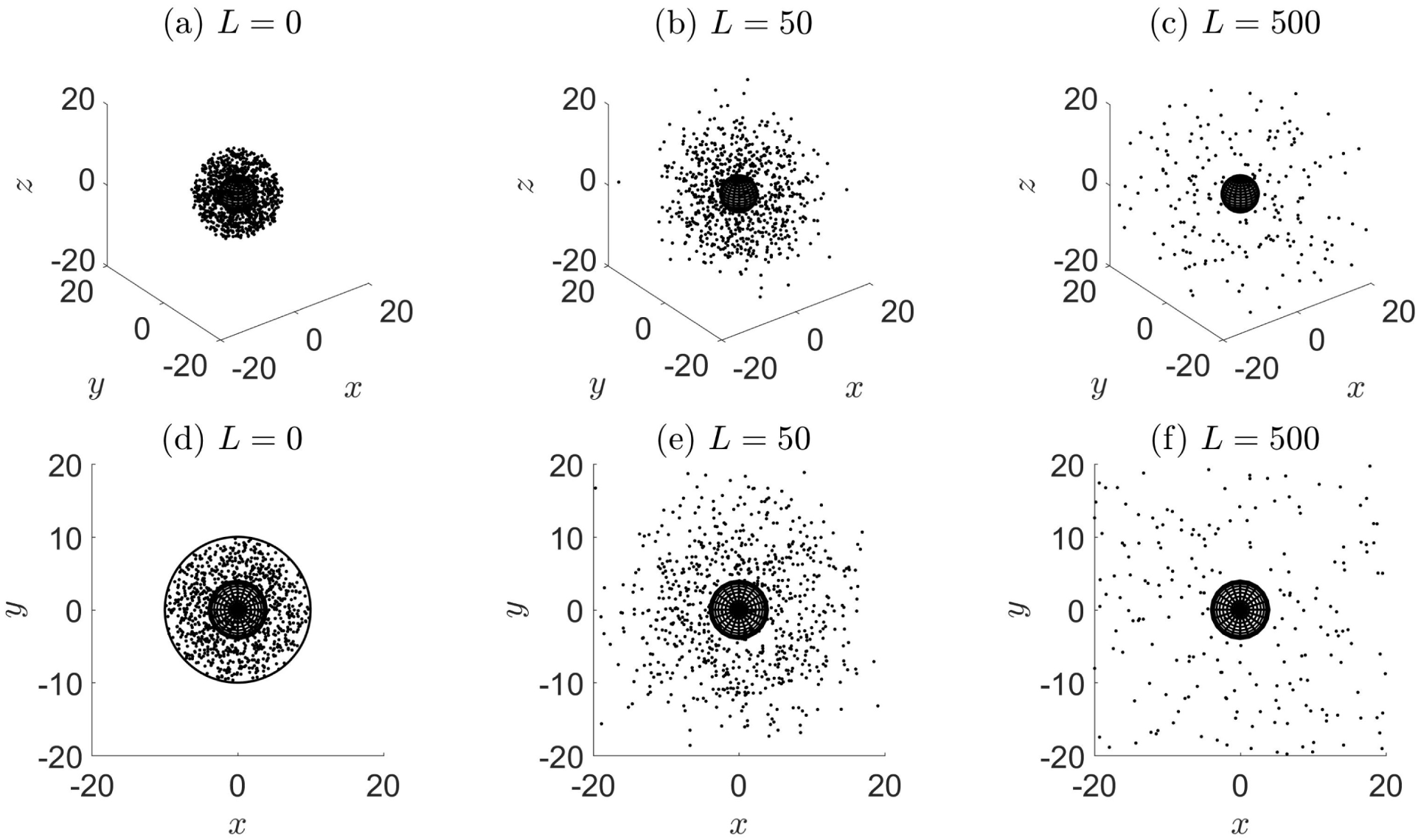
Evolution of the 3D spatial distribution for a population of *N* = 1000 individuals, uniformly distributed at (a) *L* = 0 (initial condition), within a distance *R* = 10 from the centre of the spherical trap of radius *r*_*s*_ = 4. Each individual walker performs a SRW with Gaussian increments and mobility parameter *σ* ^*^ = 1 (corresponding to sinuosity *S* = 1.79). Individual location is plotted until it is trapped or it travelled a path of maximum length (b) *L* = 50 and (c) *L* = 500, corresponding to approximately *n* = 31, 313 steps, respectively. Plots (a)-(c) present a 3D view and (d)-(f) presents a top-down view of the above. The black circle in (d) is included to illustrate that the walkers are confined within the vicinity at *L* = 0, but later move in unbounded space.

**Figure 2.3.2.**
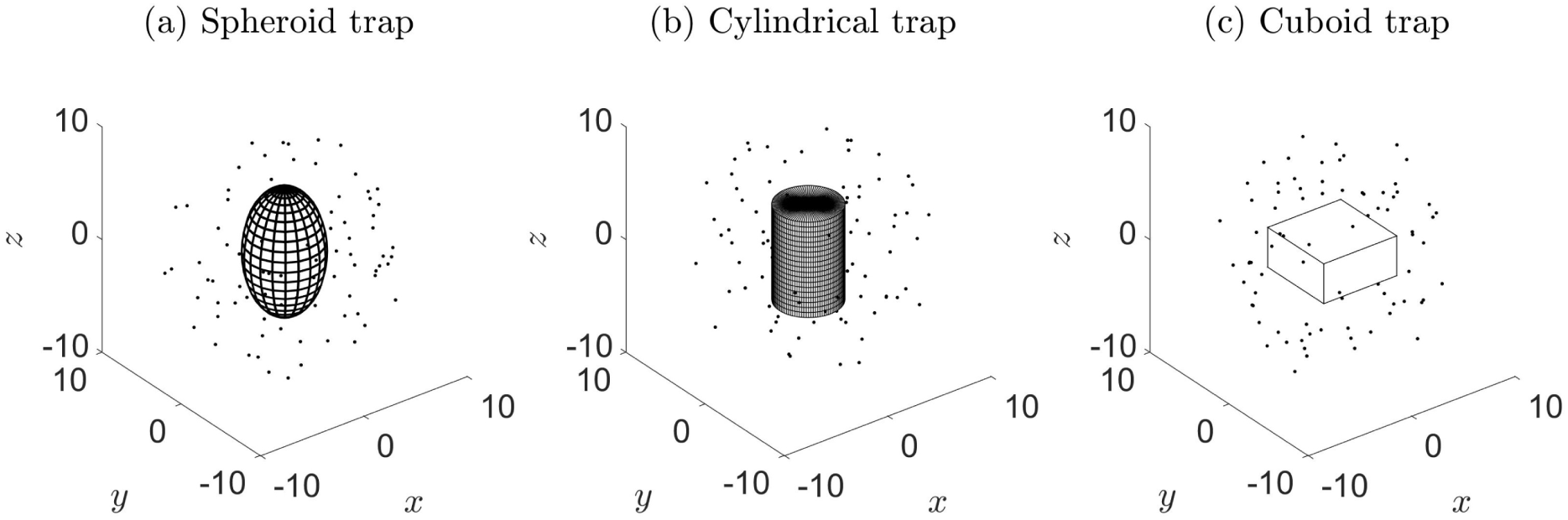
Illustration of the different trap shapes. (a) (Prolate) Spheroid trap: radius *r*_*s*_ = 3.27, *h*_*s*_ = 5.56, *ε* _*s*_ = 1.7, (b) Cylindrical trap: radius *r*_*c*_ = 2.82, *h*_*c*_ = 8.46, *ε* _*c*_ = 1.5, (c) Cuboid trap: base length *e*_*b*_ = 7.07, *h*_*b*_ = 3.54, *ε* _*b*_ = 0.5. *N* = 100 individuals are initially uniformly distributed over the vicinity between the trap and a radial distance of *R* = 10 measured from the centre of the trap. The dimensions are chosen so that the surface area, *A*, of each trap is approximately equal to 200, which is a necessary requirement to compare between these geometries (see explanation in §3.1).

with the specific case *r*_*s*_ = *h*_*s*_ reduces to a spherical shaped trap.

2. Cylindrical trap with radius *r*_*c*_ and height *h*_*c*_,

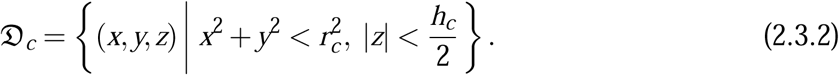

3. Rectangular cuboid trap with square base of side length *e*_*b*_ and height *h*_*b*_,

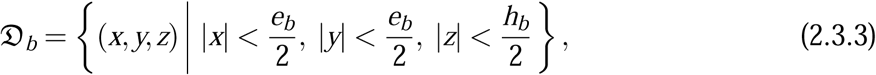

with the specific case *e*_*b*_ = *h*_*b*_ reduces to a cube shaped trap.

Subscripts ‘*s, c, b*’ refer to the spheroid, cylindrical, cuboid types, respectively.

For any trap type, we can specify its shape by introducing dimensionless elongation parameters. For the spheroid, we considered the ratio of polar to equatorial radii 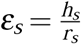, where *ε* _*s*_ < 1 corresponds to an oblate spheroid and *ε* _*s*_ > 1 to a prolate spheroid. For the cuboid we considered the ratio of height to base side length 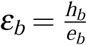, where *ε* _*b*_ = 1 corresponds to a cube, and for the cylinder we considered the ratio of height to base diameter 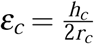.

We can then write expressions for the total surface area as:

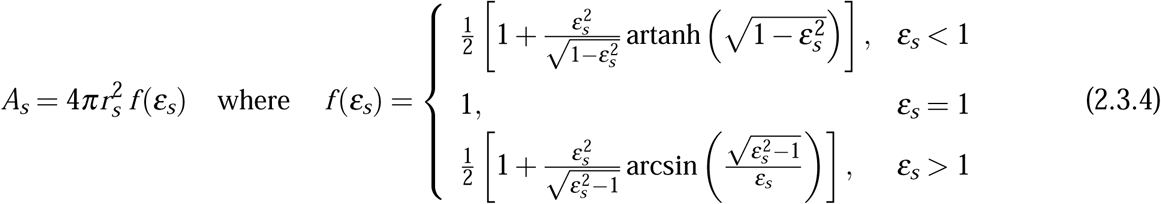

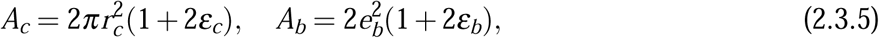

and for volume:

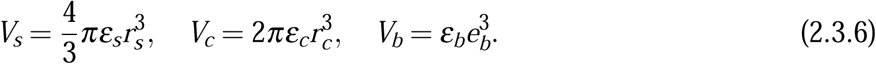

We can also express volume as a function of area as:

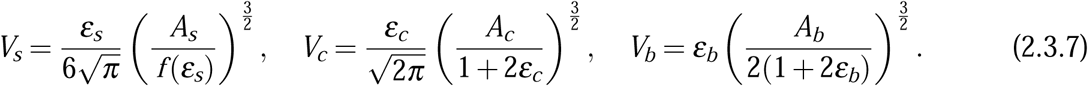

The initial distribution of individual location is considered to be uniform over a vicinity, which is defined as the space between the trap and some fixed outer distance *R*, measured from the centre of the trap. In the case of a spherical trap, we can think of this as the 3D extension of uniformly distributed points on an annulus, i.e. between two concentric spheres. If we describe initial location in spherical co-ordinates as **x**_0_ = (*r*_0_, *θ* _0_, *ϕ* _0_), then the corresponding probability density functions can be written explicitly as:

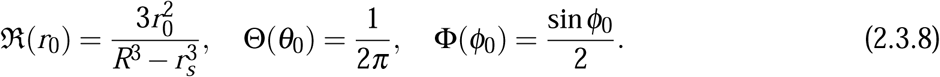

Using the inverse transform technique (Grimmet and Stirzaker, 2001), the initial location of each individual can then easily be simulated by:

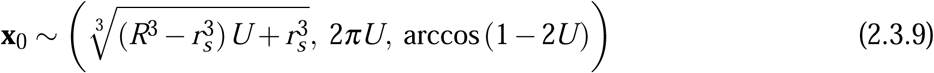

where *U* is a random variable drawn from the uniform distribution between 0 and 1.

In the case of other trap shape, the vicinity no longer has an infinite number of symmetry axes and therefore, to simulate a homogeneous population is not as straightforward. In these cases, we drawn the initial locations at random in the whole sphere of radius *R*, and removed those occurring within the trap.

## 3 Results

### 3.1 Effect of trap shape type

Consider the usual simulation setting outlined in § 2.3, for a spherical trap (*ε* _*s*_ = 1) with increasing trap size.

By simulating trap count data for different sized spherical traps, we can investigate whether captures are better correlated with trap surface area or trap volume. This approach can also be applied to cuboid and cylindrical traps.

**Figure 3.1.1.**
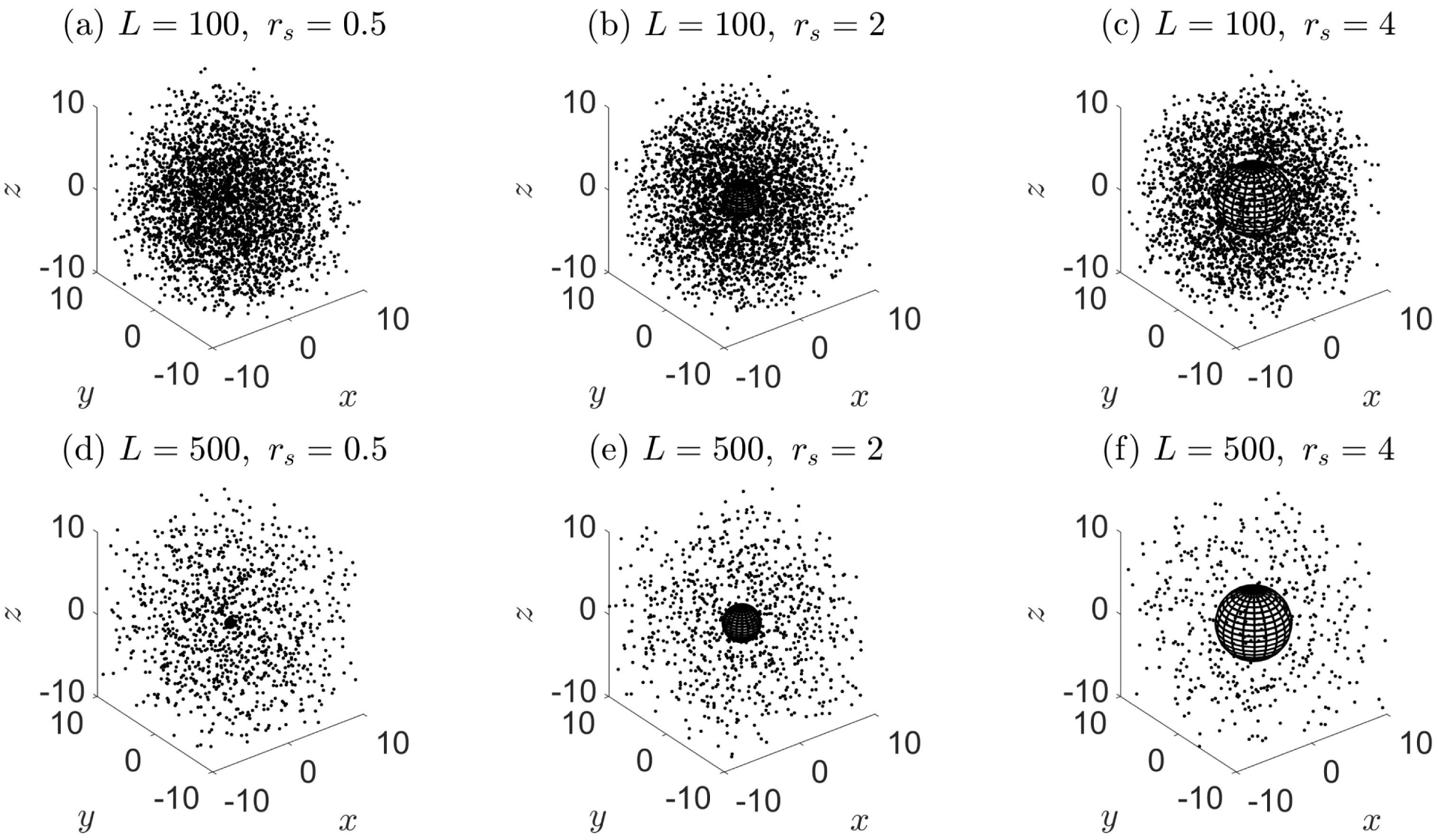
Snapshots of the spatial distribution in the case of spherical traps with radii *r*_*s*_ = 0.5, 2, 4, (surface area *A* = 3.14, 50.27, 201.06), after a maximum path length of (a)-(c) *L* = 100 and (d)-(f) *L* = 500 has been reached. Each individual executes a SRW in unbounded space with mobility parameter *σ* ^∗^ = 1 (*S* = 1.79).

We considered a cube trap *h*_*b*_ = *e*_*b*_ (*ε* _*b*_ = 1), and a ‘normalized’ cylinder where the height is equal to the base diameter *h*_*c*_ = 2*r*_*c*_ (*ε* _*c*_ = 1). The normalized cylinder and the cube lie within a sphere of radius *R* provided that the following inequalities apply:

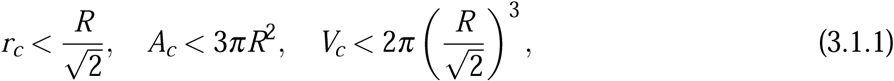

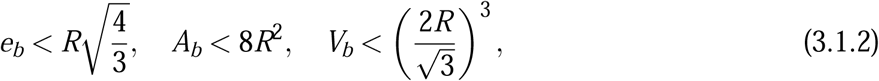

which we use to determine the range of trap dimensions, areas and volumes.

The simulated trap counts are shown in Fig. 3.1.2. It is readily seen that the cumulative trap count is a monotonously increasing, nonlinear function of trap surface area and volume. Note that the order of trap shapes, in terms of capture efficiency, is reversed depending on whether we consider the traps to have equal total area or volume.

**Figure 3.1.2.**
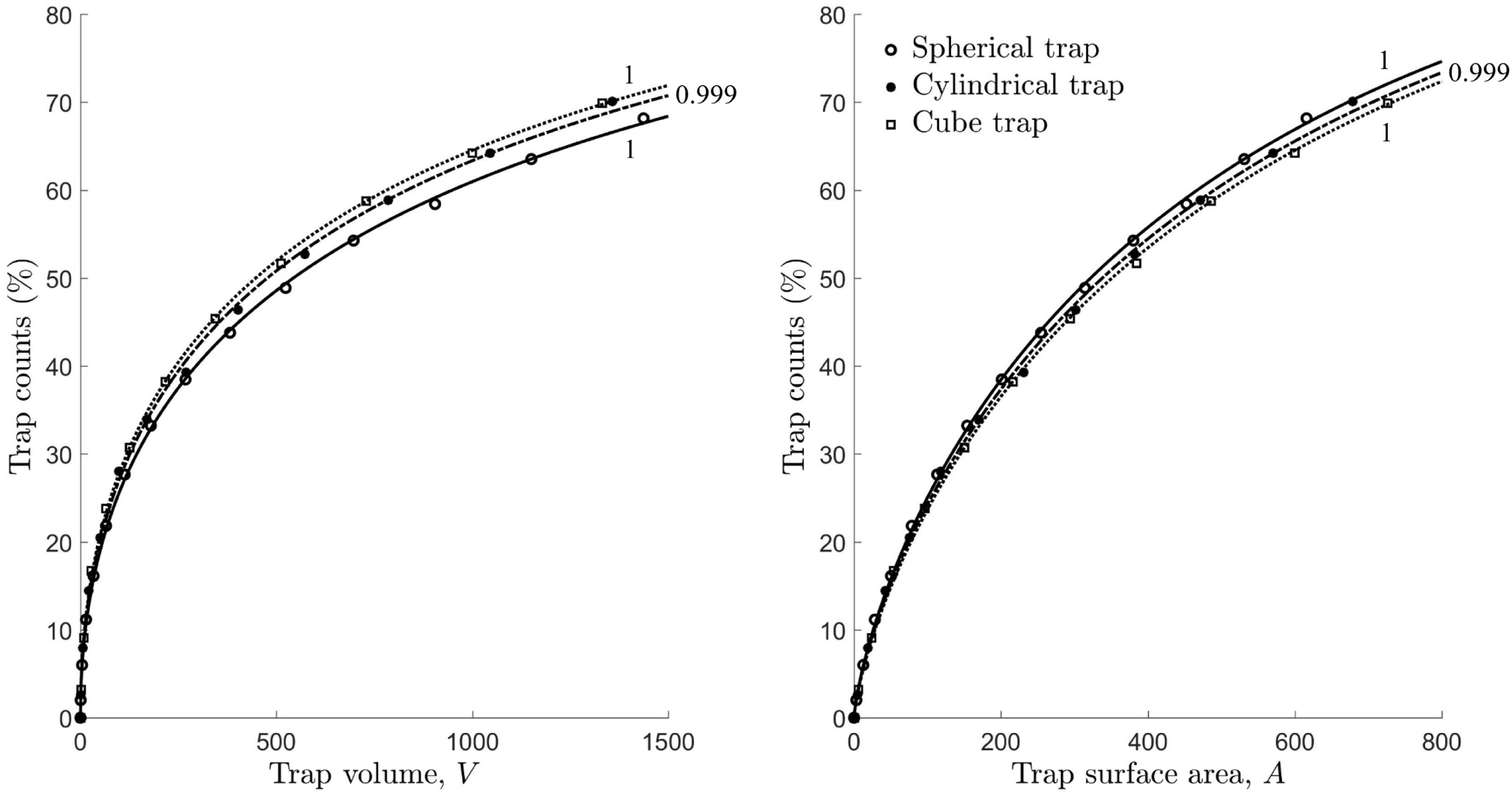
Cumulative trap counts as a function of (a) trap volume and (b) trap surface area, using non-linear regression. Solid curves are for the spherical trap, dashed-and-dotted curves are the cylindrical trap and dotted curves are for the cube trap. (a) 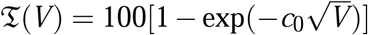 with *c*_0_ = 0.0297 for the sphere with radii *r*_*s*_ = 0.5, 1, …, 7, *c*_0_ = 0.0317 for the cylinder with radii *r*_*c*_ = 0.5, 1, …, 6 (*ε* _*c*_ = 1), and *c*_0_ = 0.0328 for the cube with base lengths *e*_*b*_ = 1, 2, …, 11 (*ε* _*b*_ = 1). (b) Same formula as in (a) with *V* expressed in terms of *A*, given by the equations in (2.3.7). The values noted alongside each curve are the squared correlation coefficients. The range of volumes/area considered are found from the upper bounds in (3.1.2). Simulation details: the movement type used is a SRW with *σ* ^∗^ = 1 (*S* = 1.79). Trap counts are recorded after a maximum path length of *L* = 500 has been reached.

### 3.2 Effect of trap elongation

In the following, we investigate the variation in trap counts for different configurations of spheroidal, cylindrical and cuboid traps, assuming the same total surface area or volume.

Trap counts for a given volume and a given trap shape (Fig. 3.2.1a) varies a lot, but the variation as a function of the elongation parameter is mainly due to a variation of area. Indeed, the sharp increase in the trap count seen in Fig, 3.2.1a for small *ε* is an immediate consequence of the fact that the decrease in *ε* to values *ε* ≪ 1 makes the shape almost flat. In order to preserve the volume, the area then becomes large. On the contrary, when the area is kept constant for all trap shape types and elongation parameter, we found that the number of captures does not vary much (Fig. 3.2.1b). In this context, spheroidal traps slightly outperform cylindrical and cuboid traps in terms of capture efficiency. As elongation has no noticeable effect (for each type of trap) whereas this factor changes the volume for a given area, it makes sense to consider traps with the same area for subsequent analyses of the possible effects of short-term persistence, long-term directional bias and diffusion of the walk.

**Figure 3.2.1.**
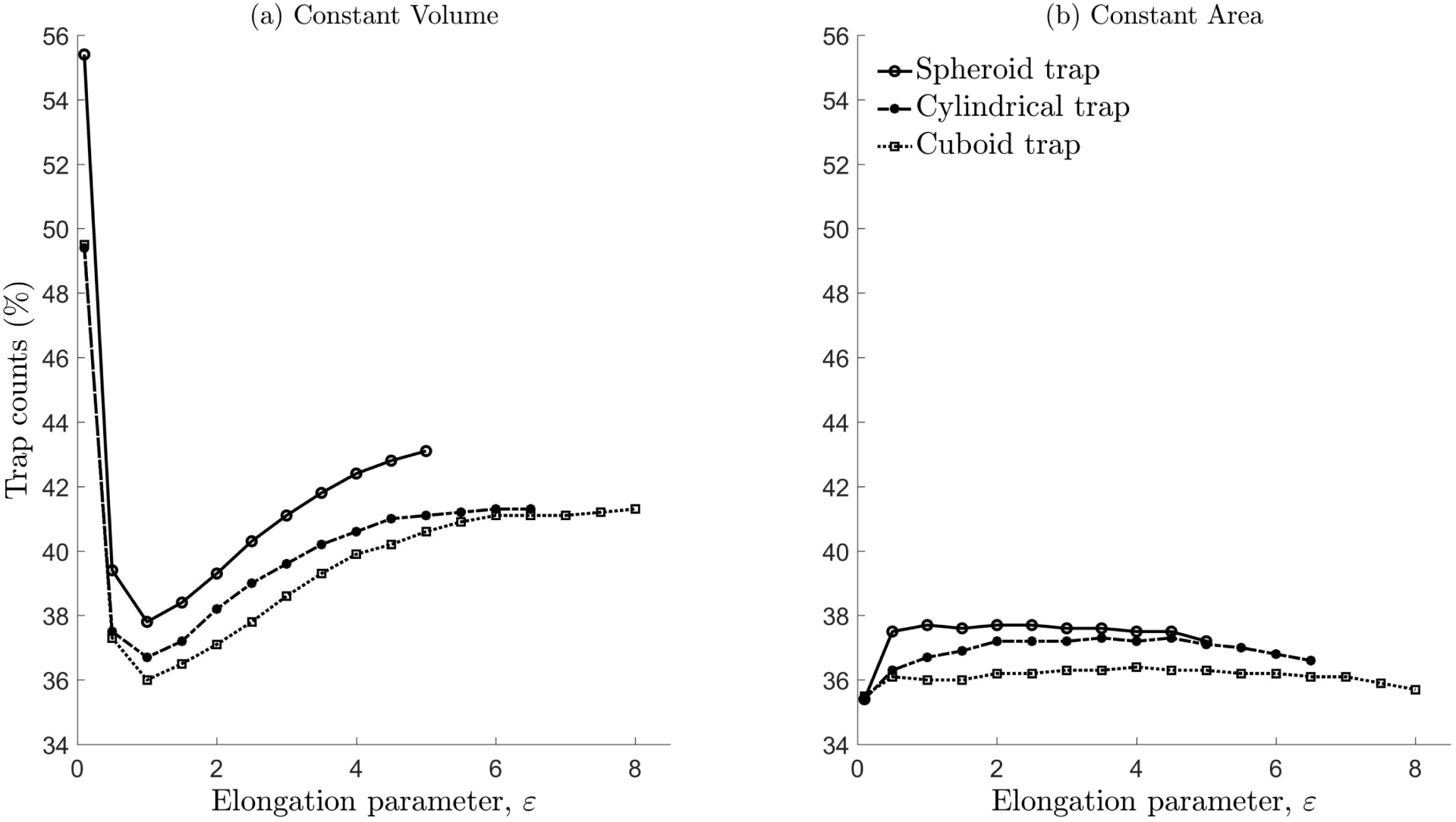
Trap captures (%) for spheroid, cylindrical and cuboid traps. (a) Each trap type has the same volume: spheroid *V* = 265.96, cylinder *V* = 217.16 and cuboid *V* = 192.45, corresponding to an area *A* = 200 for elongation parameter equal to 1. (b) All traps have the same surface area *A* = 200. The range of *ε* values considered has upper bounds *ε* _*s*_ ≤ 5, *ε* _*c*_ ≤ 6.5 and *ε* _*b*_ ≤ 8 so that all traps lie within a sphere of radius *R* = 10. The movement type considered is a SRW with *σ* ^∗^ = 1 (*S* = 1.79), and each walker is allowed to travel up to a maximum path length *L* = 500. All other details regarding the simulation setting are the same as in the caption of Fig. 3.3.1.

### 3.3 No effect of short-term persistence when diffusion is kept constant

Fig. 3.3.1 demonstrates that the inclusion of short-term persistence results in identical trap counts, assuming that all individuals perform a path with the same diffusion and same maximum path length irrespective of trap geometry.

**Figure 3.3.1.**
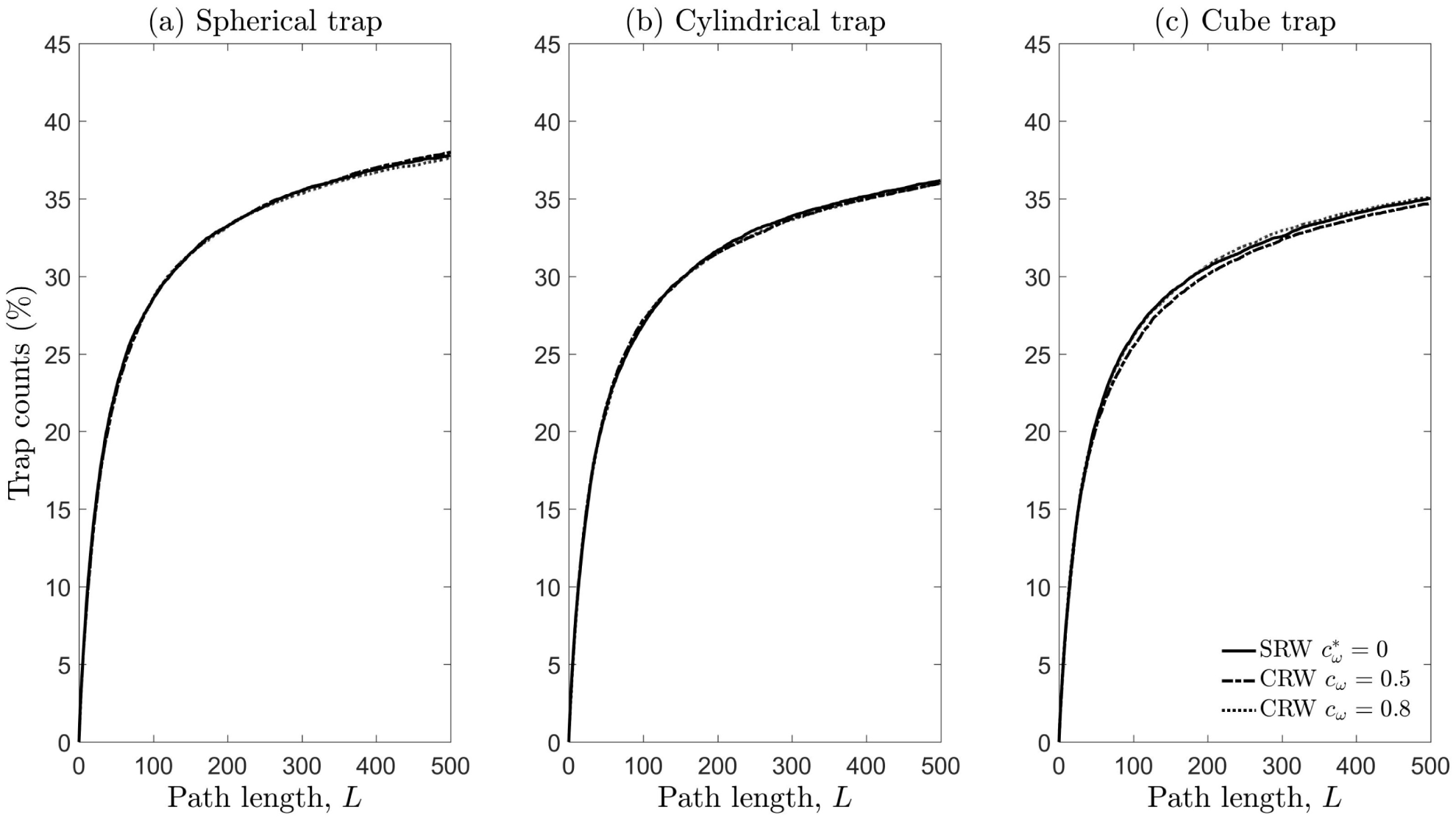
Captures (%) plotted against path length *L*. Trap geometries considered are (a) spherical *r*_*s*_ = 3.9894 (*ε* _*s*_ = 1), (b) cylindrical *r*_*c*_ = 3.2574 (*ε* _*c*_ = 1) and (c) cube *e*_*b*_ = 5.7735 (*ε* _*b*_ = 1) with equal surface area *A* = 200. Initial population is homogeneously distributed over the volume outside the trap and within a sphere of radius *R* = 10. Movement types considered are SRW with mean cosine 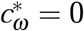, *κ* ^∗^ = 0 and *σ* ^∗^ = 1, CRW with *c*_*ω*_ = 0.5, *κ* = 1.7968, *σ* = 0.3707, and CRW with *c*_*ω*_ = 0.8, *κ* = 4.9977, *σ* = 0.1284. Scale parameters are chosen so that each movement type has the same sinuosity (*S* = 1.79) and therefore the same MSD after a large number of steps for a given path length (see equations (2.2.7) and (2.2.8)).

### 3.4 Effect of diffusion

Fig. 3.4.1 confirms that spherical traps are, on average, the most efficient. Trap counts decrease with increasing diffusion, as soon as the maximum path length is sufficiently long. We observe small but noticeable differences in efficiencies on comparing the cube and cylindrical traps. This indicates that the impact of trap geometry can be important in this case. Also, we note that trap counts accumulate much slower if diffusion is low, and given that the path length is small. This has an intuitive interpretation that individuals, on average, do not have enough time to approach the trap.

**Figure 3.4.1.**
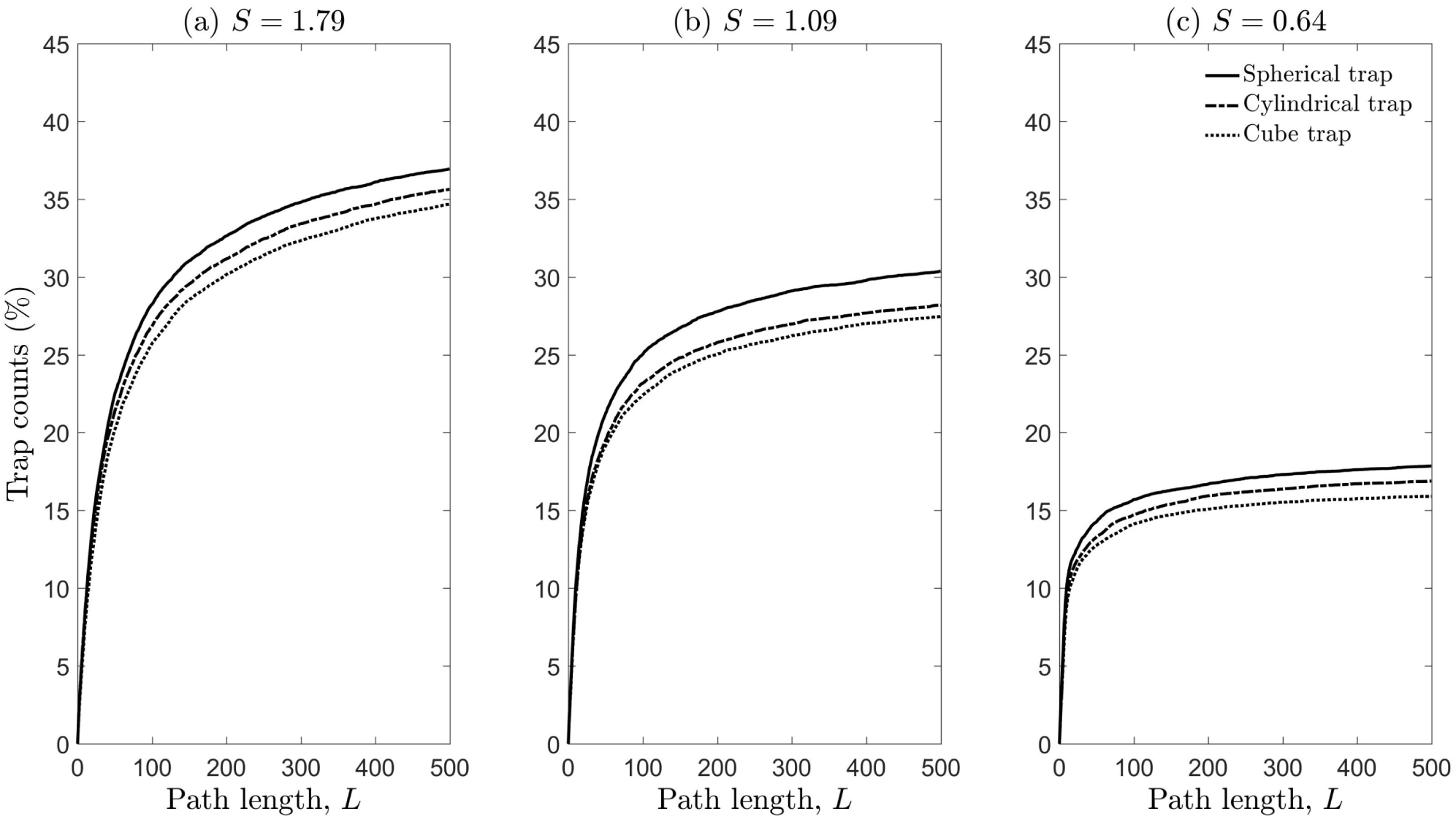
Captures (%) plotted as a function of path length for a Spherical, Cube and Cylindrical trap with mean cosines (a) 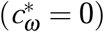, (b) *S* = 1.09 (*c*_*ω*_ = 0.5) and (c) *S* = 0.64 (*c*_*ω*_ = 0.8). Contrary to what occurs in Fig. 3.3.1, the scaling parameter was the same for all walks (*σ* ^∗^ = *σ* = 1) so that the diffusion increases with *c*_*ω*_.

Fig. 3.4.2 shows that trap counts decrease, on average, with increasing mean cosine (i.e. increasing short-term persistence/diffusion), for all trap shapes. It is worth noting that, when diffusion is large, trap count differences decrease, implying that the impact of trap geometry is then not that important. For relatively smaller values of diffusion, there is a clear hierarchy of trap shape in terms of trapping efficiency, with the spherical trap retaining the most counts, followed by the cylindrical trap, and then the cube.

**Figure 3.4.2.**
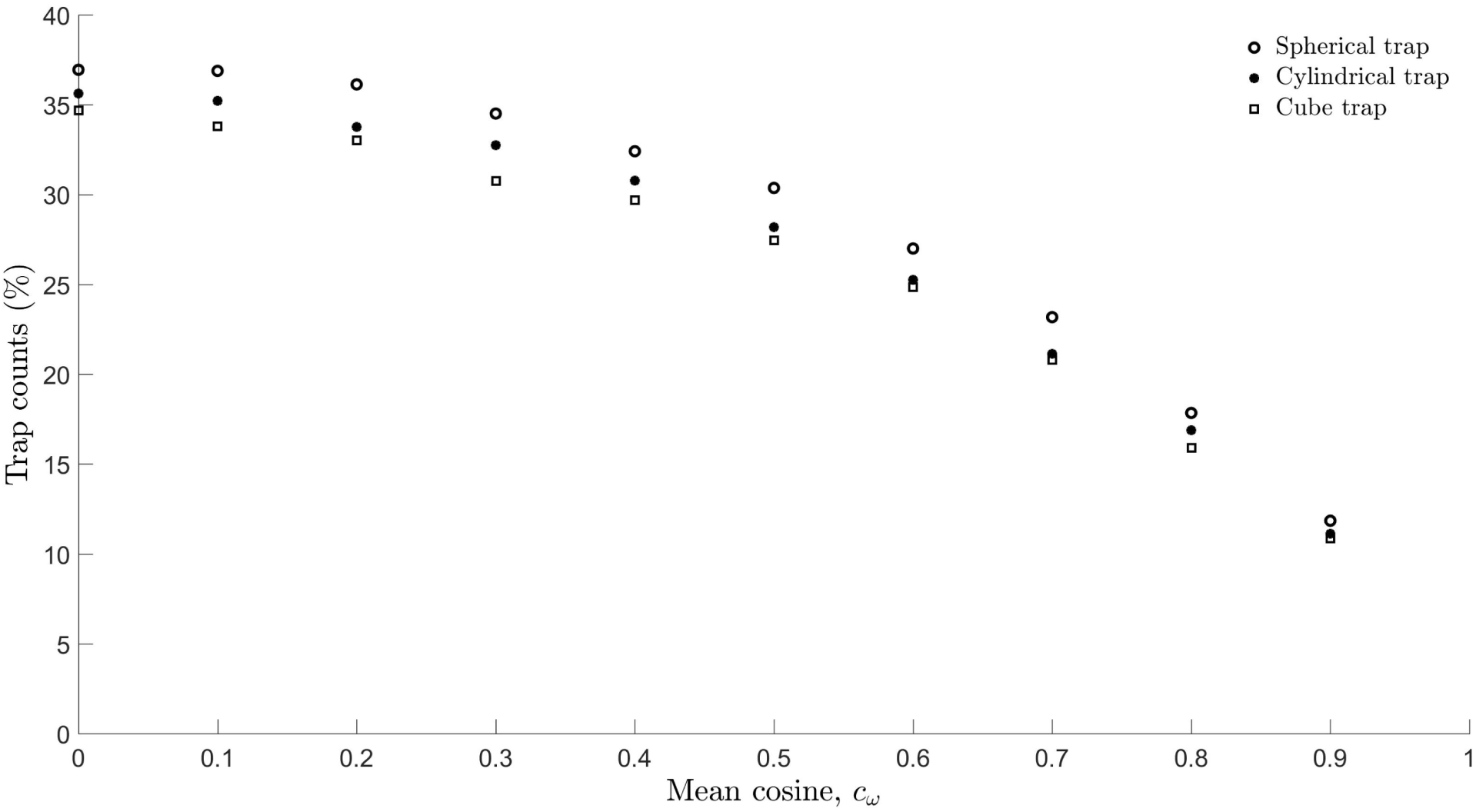
Captures (%) as a function of mean cosines for a Spherical, Cube and Cylindrical trap. Sinuosity values range from *S* = 1.79 for 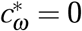 to *S* = 0.44 for *c*_*ω*_= 0.9. All details are the same as that described in the caption of Fig. 3.4.1.

### 3.5 Effect of long-term bias

Fig. 3.5.1 shows that the presence of long-term bias towards the trap, as expected, dramatically increases captures. There is a clear hierarchy of trap shapes in terms of capture efficiency.

**Figure 3.5.1.**
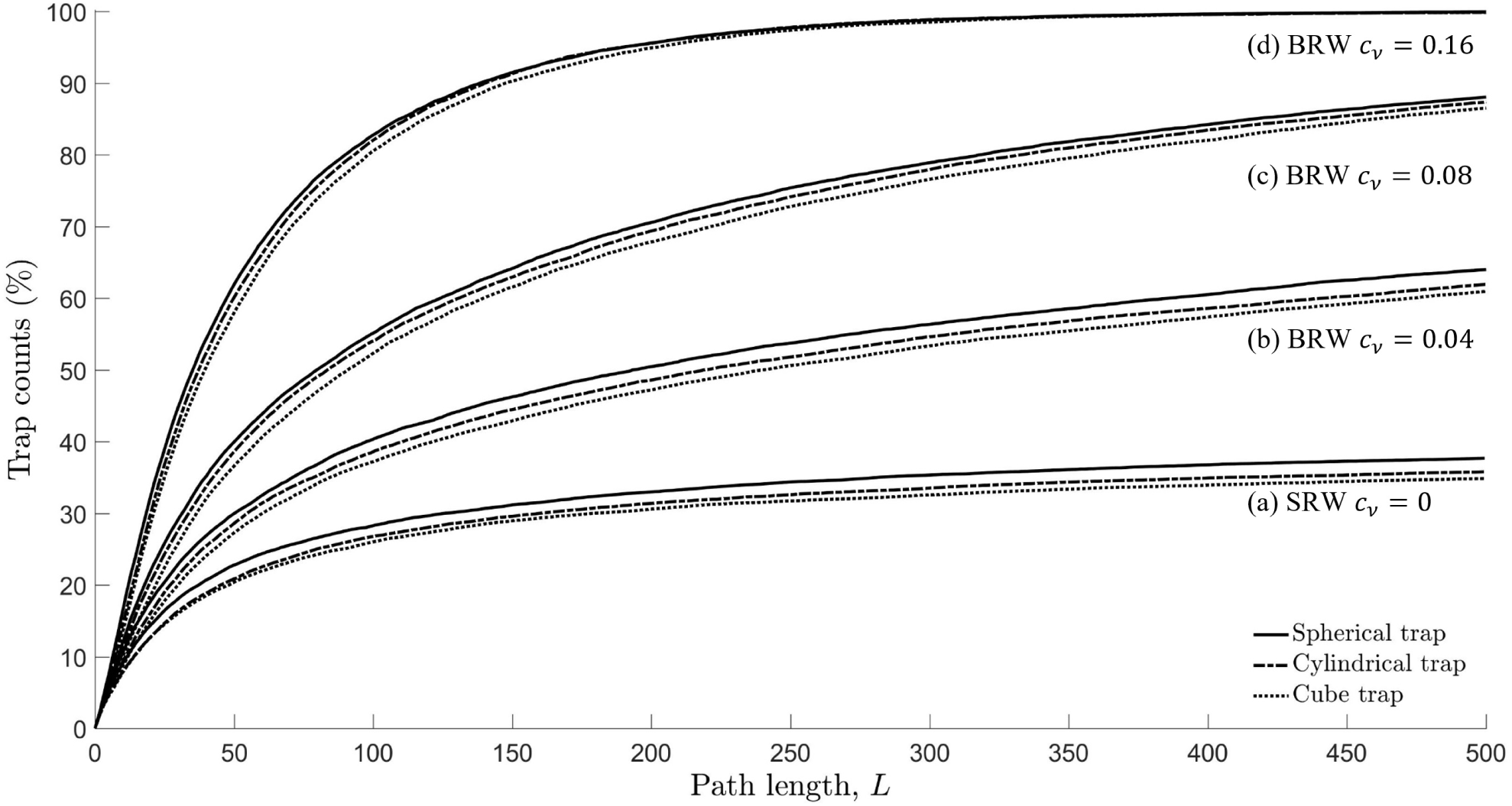
Captures (%) plotted against path length *L* for different trap types. Movement types considered: (a) SRW with mean cosine 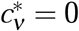 and *σ* ^∗^ = 1, (b) BRW *c*_ν_= 0.04, *κ* ′ = 0.1201, *σ* ′ = 1.0009, (c) BRW *c*_ν_= 0.08, *κ* ′ = 0.2409, *σ* ′ = 1.0038, (d) BRW *c*_ν_= 0.16, *κ* ′ = 0.4876, *σ* ′ = 1.0233. Scale parameters are chosen so that the BRW is asymptotically equivalent to a SRW in terms of diffusion. All other details, such as trap dimensions, are exactly the same as in the caption of Fig. 3.4.1.

## 4 Discussion

Dispersal and movement are fundamental for understanding the distribution and abundance of species in ecosystems. All species change their location in space at least during some stages of their life. Movement is known to have fundamental implications for individual survival, behaviours and reproduction, the population dynamics, and on fitness and evolution (Clobert et al., 2001; Bullock et al., 2002). The capacity for movement is prolific across different species. For instance, while plants do not normally move, their seeds and spores do and can cover considerable distances before settling down. Insect eggs and pupae do not move, but larvae and/or adults move most of the time, e.g. to forage for food. Most vertebrates move practically all their life, e.g. to forage, to avoid predators, to look for a mating partner, etc. Understanding of the typical movement patterns is therefore a major focus of ecology and population biology (Turchin, 1998).

Among many research tools available to study individual animal movement, mathematical modelling plays an increasingly important role (Turchin, 1998; Codling et al., 2008). Random walks (RWs) are appropriate approaches for understanding species movement patterns particularly as a stochastic or statistical description of dispersal. They are easy to implement: it is rather straightforward to investigate movement paths using computer simulations based on RWs. More importantly, by considering individual movement as a stochastic process, it is often possible to obtain a general analytical description, in terms of the dispersal kernel and/or the statistical moments, as functions of time, and thus to reveal generic properties of different movement behaviours (Reynolds, 2010; Codling and Plank, 2011; James et al., 2011; McClintock et al., 2012; Tilles and Petrovskii, 2015; Tilles et al., 2017). While there has recently been considerable progress in understanding these issues, most theoretical studies on animal movement have been predominantly limited to 2D cases. Meanwhile, in the real-world application of monitoring flying insects (e.g. different taxa of fruit flies), traps are usually elevated above the ground, sometimes at a significant height, for e.g. 1-10 metres (Epsky et al., 2004). Thus, the movement of flying insects in the vicinity of an elevated trap is essentially performed in 3D space, and hence it should be modelled as such.

Understanding the efficiency of trapping resulting from the interplay between the movement pattern (as described by the SRW, CRW and BRW) and the shape of the trap was the focus of this study. We first derived the expression for the MSD as a function of time (or number of steps), and conditions of equivalence between RWs with different step size distributions were obtained in terms of diffusion. We then proceeded to numerical simulations of trap counts with traps of different shapes commonly used in ecological studies, i.e. spheroid, cylinder and cuboid. As one result of immediate practical importance, we revealed the non-linear dependence of trap counts on the geometry of traps, quantified by either the area of the trap surface or the trap volume, and provided corresponding analytical expressions useful for trap count estimations (see Fig. 3.1.2). On considering trap elongation, we found that trap counts do not vary much given that the surface area is fixed, and that there is a clear hierarchy in terms of which traps are more efficient, with the spheroidal trap outperforming the cylindrical trap, followed by the cuboidal trap (see Fig. 3.2.1). Also, rather counter-intuitively, we showed that the short-term persistence of the individual movement (‘micro-structure’) does not have any notable effect on the trap counts when the diffusion is kept constant (see Fig. 3.3.1), and it turns out that only the ‘macro-structure’ is important (see Figs. 3.4.1 and 3.4.2).

One application of movement models arises from the needs of ecological monitoring (Greenslade, 1964; Byers, 2012; Siewers et al., 2014; Miller et al., 2015). Monitoring of invertebrates, insects in particular, is often performed by installing traps and then interpreting trap counts (catches). The latter, however, appears to be a challenging problem. It is deceptively easy to interpret the trap counts in the relative way, i.e. ‘larger count implies larger population’, but this can be misleading or simply wrong because of the interplay between the movement activity and the population density: a small population of fast moving animals can result in the same trap count as a large population of slower moving animals (cf. ‘activity-density paradigm’ (Thomas et al., 1998). An absolute interpretation of trap counts relating them to the population density in the vicinity of the trap is possible (Petrovskii et al., 2012, 2014; Ahmed and Petrovskii, 2019) but it requires a succession of several trap counts and some information about the movement pattern such as the frequency distribution of step sizes and turning angles along the path (also the distribution of different movement modes, rest time, etc., in case of more complicated movement behaviours) as well as a good understanding of the effect of trap geometry (Ahmed and Petrovskii, 2019).

Furthermore, in the statistical application of models to ecological data, a pervasive and recurrent problem is understanding the biases introduced through the measurement or observation of the ecological system (e.g. Hilborn and Mangel, 1998; de Valpine and Hastings, 2002). More accurate estimates of the number of individuals that move or are present in a given location require the use of mathematical tools. Many distance sampling methods have been developed (Buckland et al., 2015) to link observations on counts of individuals to estimates of population size. More recently Bayesian hierarchical methods (e.g Doucet et al., 2001; Bonsall et al., 2014; Kantas et al., 2015; Bonsall et al., 2020) have been developed and applied in an ecological context to approach the decomposition of error into measurement and process components. The mathematical frameworks we develop here, provide a richer set of tools to be able to relate how the biases in individual behaviours influence measurement error problems and hence provide more robust determinants of population level measures. With a more detailed understanding of the effects of different trap geometries on capturing/detecting individuals in a population will provide more robust ways in which to discern broad scale ecological patterns.

Coupled movement and dynamical models such as integro-difference approaches (Kot and Schaffer, 1986; Lutscher, 2019) have widespread application in ecology for understanding invasion speeds (e.g. Kot, 1992), Allee effects (e.g. Wang et al., 2002), climate change (e.g. Zhou and Kot, 2011) and invasive species control (e.g. Kura et al., 2019). All rely on a dispersal kernel to relate movement from one location to another (e.g. Reimer et al., 2016, 2017) and the influence this has on the population dynamics. This dispersal kernel is critical for ensuring model predictions can be accurately validated against experiments and/or observations. Our work on 3D RWs now provides a way in which to scale up from individual movement rules to generate appropriately formulated dispersal kernels. Furthermore, the individual basis to the movement and dispersal patterns provides an alternative approach to link movement and the population dynamics without recourse to simpler mean-field approaches.

A question may arise as to why one should use RWs to model explicitly hundreds or thousands of randomly moving animals rather than the corresponding mean-field mathematical description instead. If the potential ecological applications of our work is somewhat obvious, several methodological questions remain unresolved. It is well-known that, for the SRW, the dynamics of the population density distribution over space is described by the diffusion equation (Kareiva and Shigesada, 1983; Ahmed, 2015) and for the CRW, by the Telegraph equation, (e.g. see Turchin, 1998; Codling et al., 2008). However, note that the analytical solution of the diffusion equation, from which the average trap count can be calculated (see Petrovskii et al., 2012, 2014; Ahmed and Petrovskii, 2015) is, even in case of relatively simple trap shapes such as a spheroid or cylinder, only available as a Fourier series where the exponents (the eigenvalues of the corresponding boundary problem) still need to be found numerically. With the reliancy on numerical approximations and approaches, the ‘analytical’ description of trap counts is not much different from that derived from the individual based model. In the case of more realistic movement described by the CRW, the situation is actually much more complex, as the solutions of the boundary problem for the Telegraph equation in the general case are not positively defined (Tilles and Petrovskii, 2019). Simulation of trap counts using individual based models therefore provides a robust and plausible alternative to analytical approaches.

## 5 Conclusion

In conclusion, these issues notwithstanding, we have shown how different trap geometries and the 3D movement of individuals can bias trapping efficiency. Understanding how diffusion, directed movement and trap shape can affect counts, estimates and observations has critical implications for spatial ecology and for understanding the distribution and abundance of species. These individual based, geometric approaches warrant further investigation and application in problems in contemporary spatial ecology. The next natural step that we hope to see in the near future, is analyses of real flying animal movements using 3D RW models.

## Abbreviations

3D: Three-dimensional
SRW: Simple random walk
CRW: Correlated random walk
BRW: Biased random walk

## Acknowledgements

DAA is thankful to Philip Maini (Oxford, UK) for a fruitful and stimulating discussion which helped improve the manuscript.

## Author contributions

DAA and SB participated in the design of the study, conducted the simulations, analysed the results, drafted and critically revised the manuscript. MBB and SVP participated in the interpretation and discussion of results and also critically revised the manuscript. All authors gave final approval for publication and agree to be held accountable for the work performed therein.

## Funding

This work was funded by the Kuwait Foundation for the Advancement of Sciences (KFAS) [Grant number: PR1914SM-01] and the Gulf University for Science and Technology (GUST) internal seed fund [Grant Number: 187092].

## Ethical Approval and Consent to participate

Not applicable.

## Consent for publication

Not applicable.

## Availability of supporting data

Not applicable.

## Competing interests

The authors declare that they have no competing interests.

## Notes

### Competing Interest Statement

The authors have declared no competing interest.

